# Viral replication modes in single-peak fitness landscapes: a dynamical systems analysis

**DOI:** 10.1101/239921

**Authors:** Joan Forners, J. Tomás Lázaro, Tomás Alarcón, Santiago F. Elena, Josep Sardanyés

## Abstract

Positive-sense, single-stranded RNA viruses are important pathogens infecting almost all types of organisms. Experimental evidences from mutant distributions and amplification kinetics of viral RNA suggest that these pathogens may follow different RNA replication modes, ranging from the stamping machine replication (SMR) to the geometric replication (GR) modes. Despite previous theoretical works have focused on the evolutionary dynamics of RNA viruses amplifying their genomes with different strategies, few is known in terms of the bifurcations and transitions involving error thresholds (mutation-induced dominance of mutants) and lethal mutagenesis (mutation-induced extinction of all sequences). Here we analyze a dynamical system describing the intracellular amplification of viral RNA genomes evolving on a single-peak fitness landscape focusing on three cases considering neutral, deleterious, and lethal mutants spectra. In our model, the different replication modes are introduced with parameter *α*: with *α* ≳ 0 for the SMR and *α* = 1 for the GR. We analytically derive the critical mutation rates causing lethal mutagenesis and error catastrophe, governed by transcritical bifurcations that depend on parameters *α*, *k*_1_ (replicative fitness of mutants), and on the spontaneous degradation rates of the sequences, *ϵ*. For the lethal case the critical mutation rate involving lethal mutagenesis is 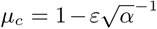. Here, the SMR involves lower critical mutation rates, being the system more robust to lethal mutagenesis replicating closer to the GR mode. This result is also found for the neutral and deleterious cases, but for these later cases lethal mutagenesis can shift to the error catastrophe once the replication mode surpasses a threshold given by 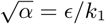.

## I. INTRODUCTION

RNA viruses are characterized for being fast replicators and reaching enormous populations sizes within infected hosts. However, virus' fast replication comes with the cost of extremely high mutation rates due to the lack of correction mechanisms of their RNA-dependent RNA polymerases (RdRp) [1, 2]. Indeed, mutation rates are so high that viral populations are thought to replicate close to the so-called error threshold, beyond which it is not possible to retain genetic information as mutant genomes outcompete the mutation-free genome [3]. These mutation rates are orders of magnitude higher than those characteristic for their cellular hosts. While the combination of fast replication, large population size and high mutation rate create the potential for quick adaptation to new environmental conditions (*e.g*., changes in host species or the addition of an antiviral drug), in a stable environment such strategy has the drawback of generating a high load of deleterious mutations. Therefore, natural selection may have favored life history traits that may balance for the accumulation of deleterious mutations.

One of such life history traits that has received a good deal of attention is the mechanism of within-cell viral replication. In the continuous of possible modes of replication, the two extremes have been particularly well studied. At the one extreme, the stamping machine mode [4], hereafter referred as SMR, implies that the first infecting genome is transcribed into a reduced number of molecules of opposite polarity that will then be used as templates to generate the entire progeny of genomes. At the other extreme, the geometric replication mode [5], hereafter named as GR, means that the newly generated progeny also serves as template to produce new opposite polarity molecules that, themselves, will also serve to generate new progeny genomes, repeating the cycle until cellular resources are exhausted and replication ends. The actual mode of replication of a given virus may lie between these two extremes. Some RNA viruses such as bacteriophages *ϕ*6 [6] and Q*β* [7] and turnip mosaic virus [8] tend to replicate closer to the SMR. In contrast, for other RNA viruses such as poliovirus [9] or vesicular stomatitis virus [10], replication involves multiple rounds of copying per cell, and thus a mode of replication that should be closer to the GR. For DNA viruses, GR is the most likely mechanism of replication given their double-stranded nature, *e.g*., bacteriophage T2 [5]. Exception maybe be single-stranded DNA viruses, such as bacteriophage *ϕX*174, that replicates via the SMR mode because it uses a rolling circle mechanism [11].

At which point of the continuous between these two extreme modes of genome replication is a virus has important evolutionary consequences. Under SMR the frequency of mutants per infected cell follows expression 1 − *e*^−^*^µ^*, being *µ* the genomic mutation rate. Under this mode of replication, the distribution of mutants per infected cell before the action of selection will follow a Poisson distribution. However, under GR this frequency is also determined by the number of replication cycles, *k*, as mutants become amplified every time they serve as template: 1 − *e^−kμ^*. The resulting distribution of mutants for GR follows the Luria-Delbruck distribution. Taken together, these observations suggest that a given RdRp will produce more mutations per cell if the mode of replication is closer to GR than if it is closer to SMR. For this reason, it has been suggested that the SMR model has been selectively favored in RNA viruses because it compensates for the extremely high error rate of their RdRps [12–14]. However, it remains unknown whether a given virus can modify its replication mode in response to specific selective pressures to promote or down-regulate mutational output.

Despite some previous theoretical work aiming to explore the implications of the different modes of replication, the evolutionary dynamics tied to both the SMR or the GR modes are not fully understood, specially the role of the topography of the underlying fitness landscape on error thresholds and, especially, on lethal mutagenesis which, to the extend of our knowledge, have not been investigated in RNA viruses with asymmetric replication modes. Indeed, the nature of the bifurcations responsible for the error threshold and lethal mutagenesis for this type of system remains unexplored. In this sense, few works have explored the effect of the mode of replication on the population dynamics of viral genomes from a dynamical point of view [15]. Recently, the analysis of a dynamical system given by a model with two variables identified a transcritical bifurcation at crossing a bifurcation threshold. For this model, the bifurcation could be either achieved by tuning the parameter that adjusted for the mode of replication or by increasing the degradation rate of the strands [16]. However, this model only considered the amplification dynamics of both (+) and (−) sense RNA strands. That is, evolution was not considered in the model.

In this article, we sought to investigate a quasispecies-like model given by a dynamical system which considers the processes of replication and mutation together with an asymmetry parameter that determines the mode by which viral RNA genomes are amplified. This parameter allows to investigate the impact of different modes of replication (either SMR, GR, or a combination of both modes of replication, see Fig. 1a). The dynamics is assumed to take place on a single-peak fitness landscape (see Fig. 1b). This landscape, albeit being an oversimplification of reality, has been widely investigated [17, 18] since it allows to group together the entire mutant spectrum in an average sequence with a lower or equal fitness than the mutation-free (master) sequence, which is located at the top of the only peak in the landscape. Such a landscape allows to consider the three different cases for the mutant sequences, given by a pool of (1) neutral, (2) deleterious and (3) lethal mutants. This dynamical system is investigated analytically and numerically focusing on three main parameters, given by: mutation rates, the mode of replication, and the fitness of the mutant sequences which allow us to consider three different mutational fitness effects mentioned above.

**FIG. 1:**
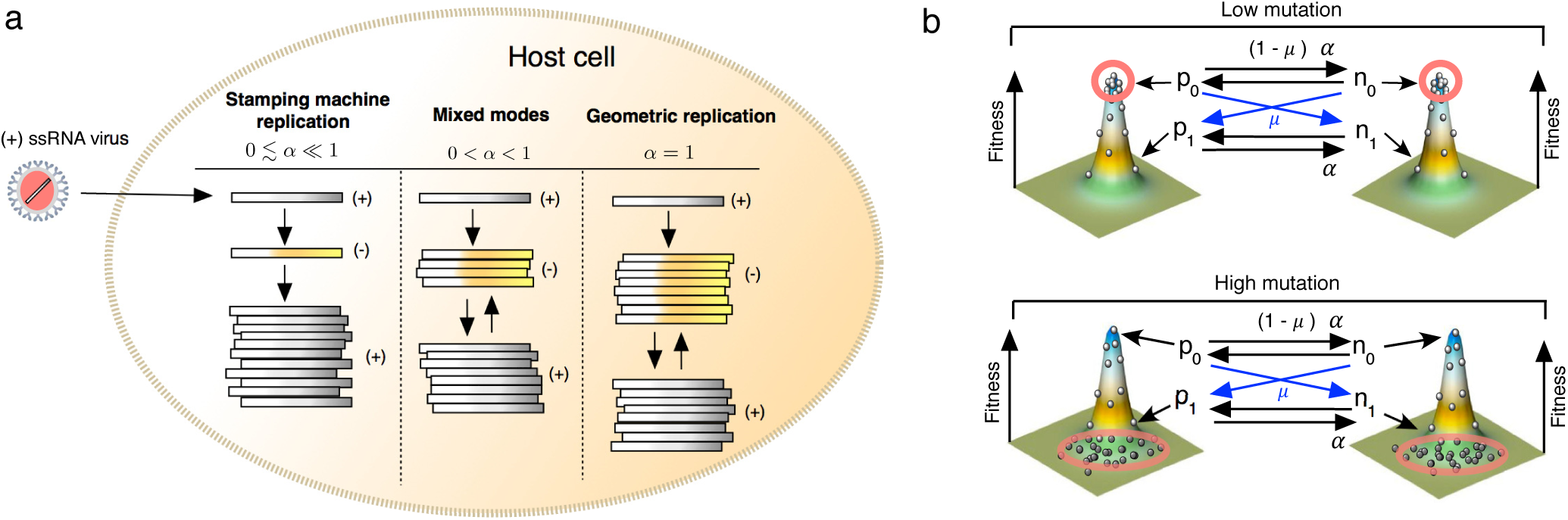
(a) Schematic diagram of the processes modeled by Eqs. (1)-(4), which consider (+) and (−) sense viral genomes (denoted by variables *p* and *n*, respectively). Upon infection, the viral genome is released within the host cell. Such a genome can be amplified following the Stamping Machine Replication (SMR) mode, the Geometric Replication (GR) model, or mixed modes. Asymmetries in replication are introduced through parameter *α* (studied as 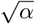): with 0 ≲ *α* ≪ 1 for SMR modes; 0 < *α* < 1 for mixed modes; and *α* = 1 for GR. (b) The model includes evolution on a single-peak fitness landscape with master (*p*_0_*,n*_0_) and mutant (*p*_1_*,n*_1_) genomes. At low mutation, the quasispecies is located at the peak, but at high mutations the quasispecies can suffer an error catastrophe and the population falls to the valley.

The structure of the paper is as follows. In Section II we introduce the mathematical model. Equilibria for this model are computed in Section III, while their stability is analyzed in Section IV. In Section V we describe the bifurcations found in the model. Section VI is devoted to some conclusions. In the Appendix Section we provide the proofs for the propositions developed in Sections III and IV.

## II. MATHEMATICAL MODEL

Here we introduce a minimal model describing the dynamics of symmetric and differential replication modes between (+) and (−) RNA viral genomes. As a difference from the model investigated in [13], which considered a more detailed description of the intracellular amplification kinetics, our model only considers the processes of replication and mutation, together with the degradation of RNA strands and their competition. The model considers four state variables, given by two classes of (+) and (−) sense viral genomes, labeled as *p* and *n*, respectively, either master (subindex 0) and mutant (subindex 1) types (see Fig. 1). The dynamical equations are defined by:

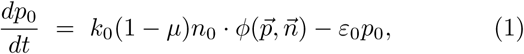

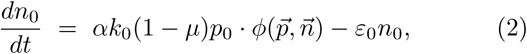

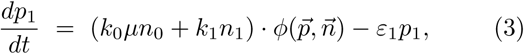

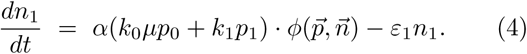

The concentration variables or population numbers span the 4*th*-dimensional open space:

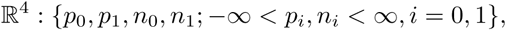
only part of which is biologically meaningful:

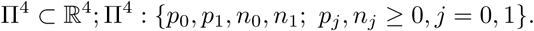

The constants *k*_0_ > 0 and *k*_1_ ≥ 0 are the replication rates of the master and the mutant genomes, respectively. Mutation rate is denoted by 0 ≤ *µ* ≤ 1. Since we are studying deleterious fitness landscapes and lethality, we will set *k*_0_ = 1. The term *ϕ*, present in all of the equations, is a logistic-like constraint, which introduces competition between the viral genomes and bounds the growth of the system [16]. This term is given by

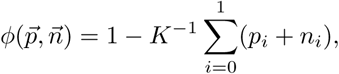

*K* being the carrying capacity (hereafter we assume *K* = 1). Parameters *ε*_0_ and *ε*_1_ correspond to the spontaneous degradation rates of master and mutant genomes, with 0 < *ε*_0_,_1_ ≪ 1. Finally, constant *α* is the parameter that introduces the mode of replication for the RNAs [16]. Two extreme cases can be identified: when *α* = 1, both (+) and (−) sense strands replicate at the same rates, following GR that results in initial exponential growth phases (see Fig. 2) [13]. When 0 ≲ *α* ≪ 1, the contribution from positive to negative strands is much lower, and thus the progeny of genomes is mainly synthesized from (-) sense genomes, giving place to SMR mode. The initial replication dynamics for the SMR replication follows sub-exponential growth (Fig. 2). Between these two extremes, our model considers a continuum of asymmetric replication modes i.e., 0 < *α* < 1.

**FIG. 2:**
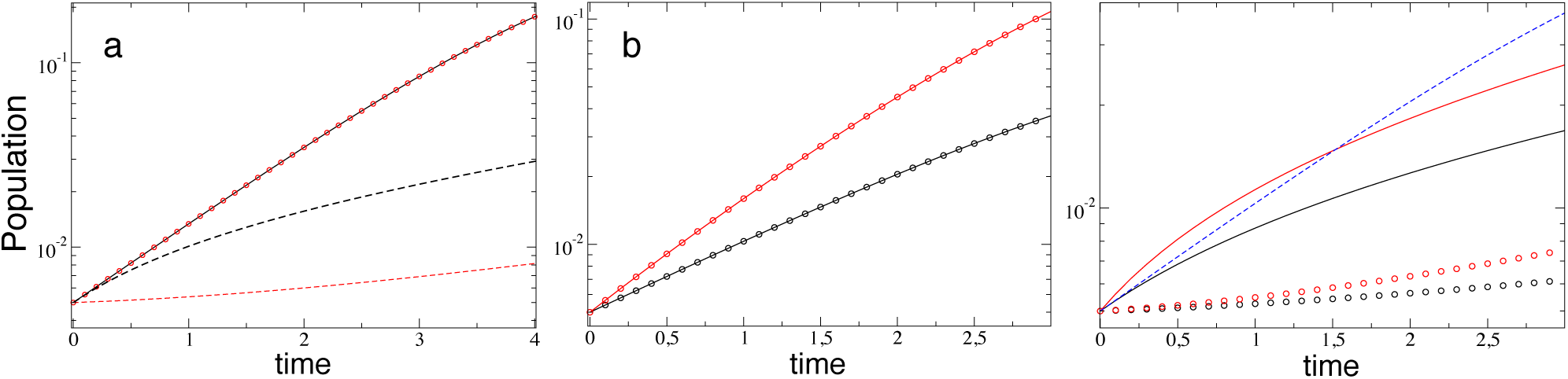
(a) Strands’ initial dynamics with *µ* = 0 and *p*_0_(0) = *n*_0_(0) = 0.005. The growth for the GR mode (*α* = 1) is exponential, resulting in a straight line in a linear-log scale: here *p*_0_ (solid black line) and *n*_0_ (red circles). The two curves below, which follow sub-exponential growth, correspond to the SMR with *α* = 0.05: *p*_0_ (dashed black) and *n*_0_ (red dashed). (b-c) Initial amplification phase with *µ* = 0.25 and *p*_0,1_(0) = *n*_0,1_(0) = 0.005. In (b) we show the dynamics for GR with *α* = 1: *p*_0_ (black solid); *p*_1_ (black circles); *n*_0_ (red solid); and *n*_1_ (red circles). In (c) we display the same results of (b) but considering SMR with *α* = 0.05. For comparison, the blue dashed line corresponds to the growth of *p*_0_ with *α* = 1 shown in (b), which results in a straight line. In all panels we set: *k*_0,1_ = 1 and *ε*_0,1_ = 10^−5^.

To simplify the exposition, we will assume the following non-restrictive assumptions on our model: (H1) equal degradation rates *ε*_0_ = *ε*_1_ = *ε* and, as mentioned, a fixed fitness value for the master genomes, setting *k*_0_ = 1; (H2) the degradation rate *ε* is smaller than the mutation rates between positive and negative strands of the master and the mutant genomes, that is, 0 < *ε* ≤ min {1 − *μ*, *k*_1_}. This last assumption involves that the dynamics is dominated by the amplification of the viral genomes, considering that degradation rates are small.

Our model assumes no backward mutations, that is, mutant sequences can not give place to master sequences. The length of RNA viral genomes (about 10^6^ nucleotides) makes the probability of backward mutations to be extremely low. This is a common assumption in quasispecies models that simplifies the dynamical equations (see *e.g*., [17–19]).

The quasispecies studied here inhabits a single-peak fitness landscape (FIg. 1b). Different heights of this fitness landscape can be studied by tuning 0 ≤ *k*_1_ ≤ 1, considering different mutational fitness effects. It is known that mutations can be deleterious, neutral, lethal, or beneficial. Some quantitative descriptions of the fitness effects of mutations reveal that about 40% of mutations are lethal, and about 20% are either deleterious or neutral. For the within-cell replication time-scale, beneficial mutations were produced with a very low percentage i.e., about 4% (see [20, 21] and references therein). Specifically, in our model we will distinguish three different cases:

1. *Neutral mutants* (*k*_0_ = *k*_1_ = 1). Mutations are neutral and thus mutant genomes have the same fitness than the master ones.
2. *Deleterious mutants* (0 ≲ *k*_1_ *<k*_0_ = 1). This case corresponds to the classical single-peak fitness landscape (see Fig. 1b), where mutations are deleterious and thus the quasispecies can be separated into two classes: the master genome and an average sequence containing all mutant sequences with lower fitness.
3. *Lethal mutants* (*k*_1_ = 0). For this case, mutations are assumed to produce non-viable, lethal genotypes which can not replicate.

## III. EQUILIBRIUM STATES

First of all, let us compute the equilibrium points of Eqs. (1)-(4) and characterize their existence conditions. That is, under which parameter values the fixed points live at the boundaries or inside the phase space Π. Let us define the following constants, which will appear in the equilibrium states (see Proposition 1) and also in their stability discussion

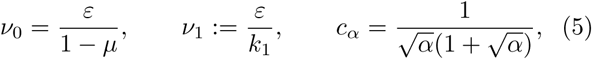
and

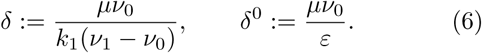

From these definitions, one has the equivalences:

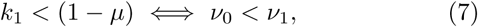

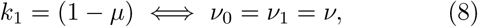

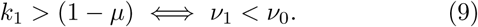

Moreover hypothesis (H2) implies that 0 < *ν*_0_ ≤ 1 and 0 < *ν*_1_ ≤ 1.

### Proposition 1

*System* (1) *presents the following equilibria:*

1. *In the Deleterious* (0 < *k*_1_ < 1) *and neutral* (*k*_1_ = 1) *cases, there are three possible equilibrium points:*

- *Total extinction: the origin*, 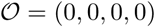
- *Master sequences’ extinction: if* 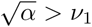 *one has the point* 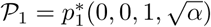, *where* 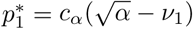.
- *Coexistence of genomes: if* 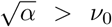 *and* 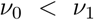, *we have* 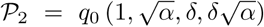, *where* 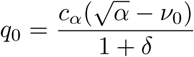.
2. *Lethal case* (*k*_1_ = 0)*. We have two equilibrium states:*

- *Total extinction: the origin*, 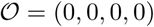
- *Coexistence of genomes: if* 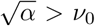 *we have the point* 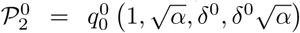 *where* 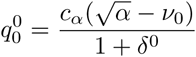.

Note that for the lethal case no equilibrium state corresponding to an error threshold is found, and only lethal mutagenesis is the alternative state to the persistence of all sequences. Figure 3 displays a diagram with the existence of the different equilibria in terms of the values of 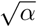 and the parameters *ν*_0_, *ν*_1_.

**FIG. 3:**
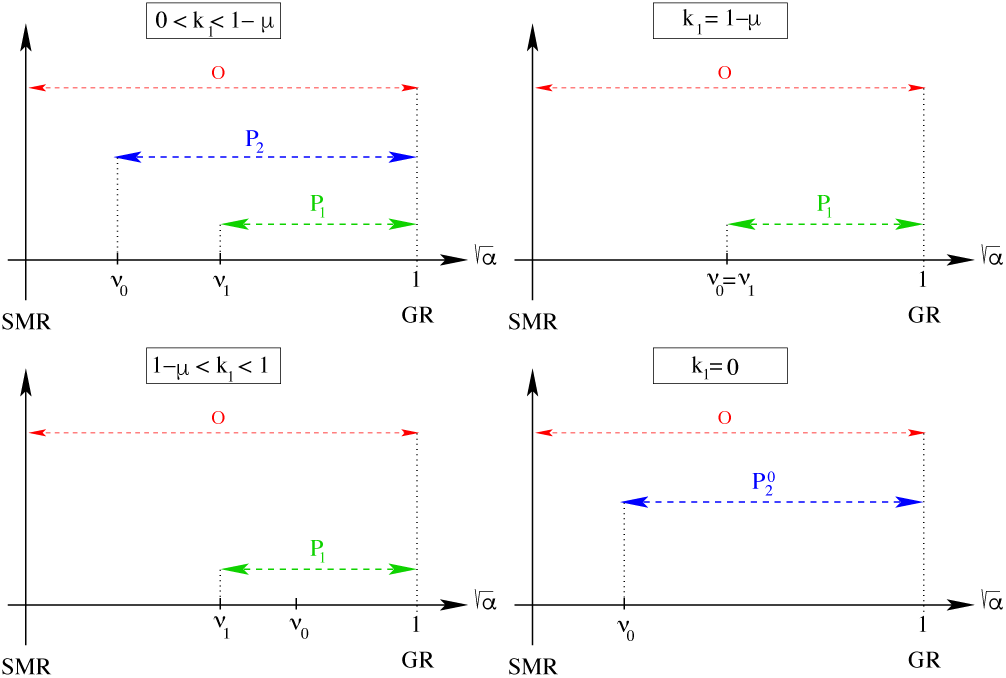
Existence of equilibria in four different scenarios: (deleterious and neutral) 0 < *k*_1_ < 1−*µ*, *k*_1_ = 1−*µ*, *k*_1_ ≥ 1−*µ* and (lethal) *k*_1_ = 0, respectively. The result are displayed increasing 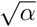 from the SMR model, with 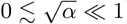) to the GR, with 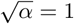) models. Here *ν*_0_ = *ε*/(1−*µ*) and *ν*_1_ = *ε*/*k*.

### Remark 1

*The coexistence points* 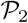 *and* 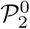 *are located on straight lines passing through the origin and director vectors* 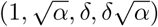 *and* 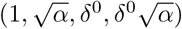.

In the case *µ* = 1, there are no master sequences *p*_0_ ↔ *n*_0_, since all master sequences mutate with probability 1. For this case, the equilibria are:

### Proposition 2

*If µ* = 1, *system* (1) *presents the following equilibria:*

1. *In the deleterious and neutral cases: the origin* 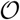 *(for any value of* 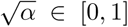*) and the point* 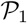 *given at the Proposition 1 provided* 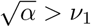.
2. *In the lethal case, the unique equilibrium is the origin* 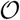, *for any value of* 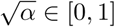.

Figure 4 displays time series achieving the equilibrium points previously described. For low mutation rates, both (+) and (−) sense strands persist, and thus 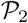 is stable (Fig. 4a). Note that close to the SMR the relative frequency of (+) and (−) strands is asymmetric, as expected, while for GR both polarities achieve similar population values at equilibrium (see also Fig. 2). The increase in mutation rates can involve the entry into error thresholds (since 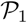 becomes stable), and the quasispecies is dominated by the mutant sequences (Fig. 4b with *α* = 0.1 and Fig. 4c for *α* = 0.1 and *α* = 0.9). The relative population of master (green) and mutant (blue) (+) sense sequences is displayed in the second and fourth columns of Fig. 4. Here also the relative frequencies of *p*_0_,_1_ achieve values close to 0.5 for the GR model, indicating that the production of both strands polarities occurs at similar rates.

**FIG. 4:**
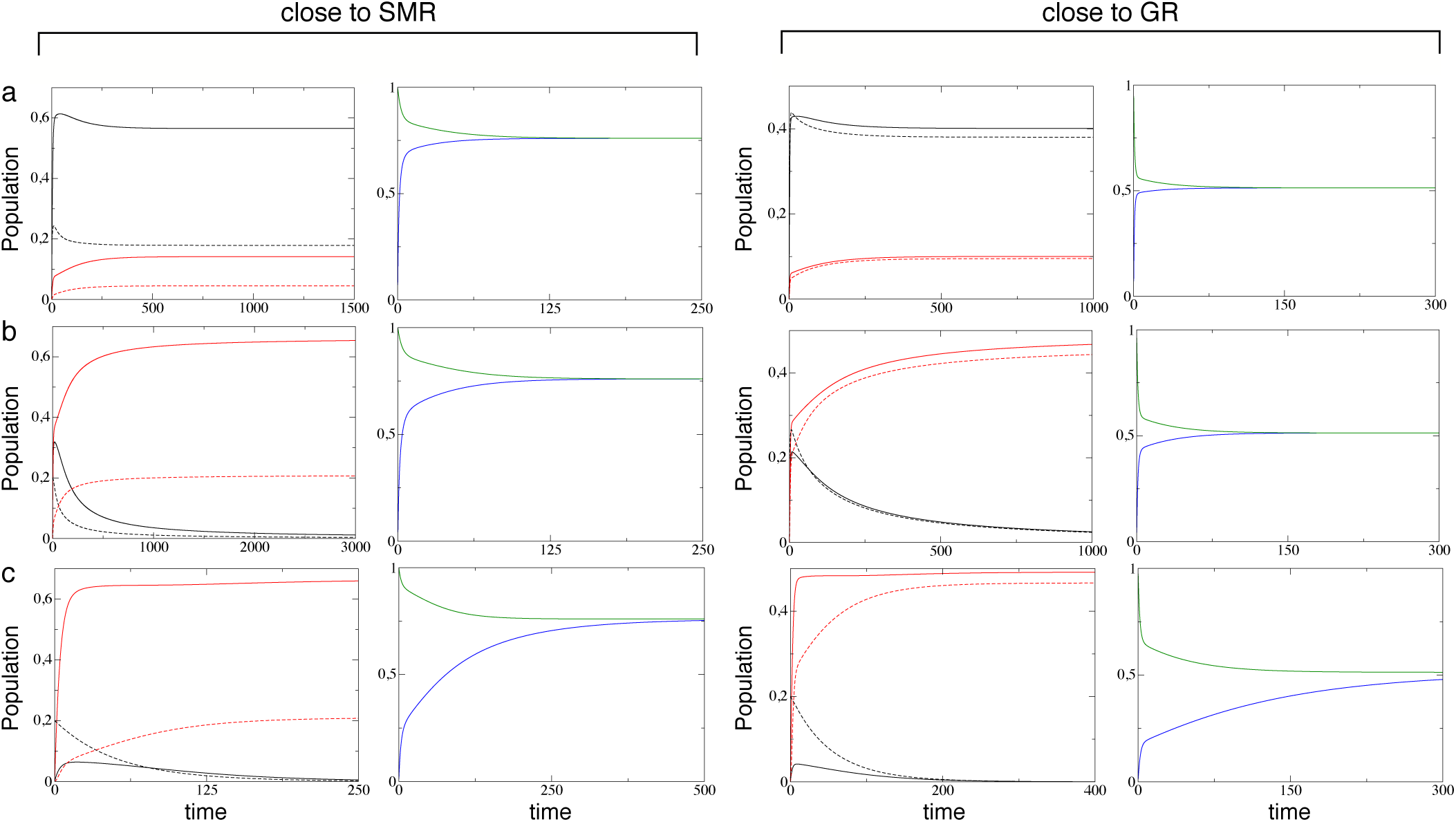
Time series for positive-(solid lines) and negative-sense (dashed lines) sequences close to the SMR (with *α* = 0.1) and close to the GR (with *α* = 0.9) modes. Here master and mutant sequences are represented in black and red, respectively. For each mode of replication: (a) *k*_1_ < (1 − *µ*) with *µ* = 0.1; (b) *k*_1_ = (1 − *µ*) with *µ* = 0.5 and (c) *k*_1_ > (1 − *µ*) with *µ* = 0.9. In all of the panels we have set *k*_1_ = 0.5, *ε* = 0.02. We also display the time series gathering the variables as follows: *p*_0_(*t*)/(*n*_0_(*t*) + *p*_0_(*t*)) (green); and *p*_1_(*t*)/(*n*_1_(*t*) + *p*_1_(*t*)) (blue).

Figure 5 displays the equilibrium populations of the four state variables at increasing mutation rates computed numerically. The first column displays the results for the SMR model (*α* = 0.1) while the second one displays the same results for *α* = 0.9, a case close to the GR model. When the fitness of the mutants is low, the SMR is less robust to extinction i.e., lethal mutagenesis, at increasing mutation, and extinction under GR takes place at higher mutation rates (see Fig. 5a). For the cases in which the fitness of mutants is higher (Fig. 5b,c) the full extinction of genomes is replaced by an error threshold, since there exists a critical value of *µ* involving the dominance of the mutant genomes and the extinction of the master sequences.

**FIG. 5:**
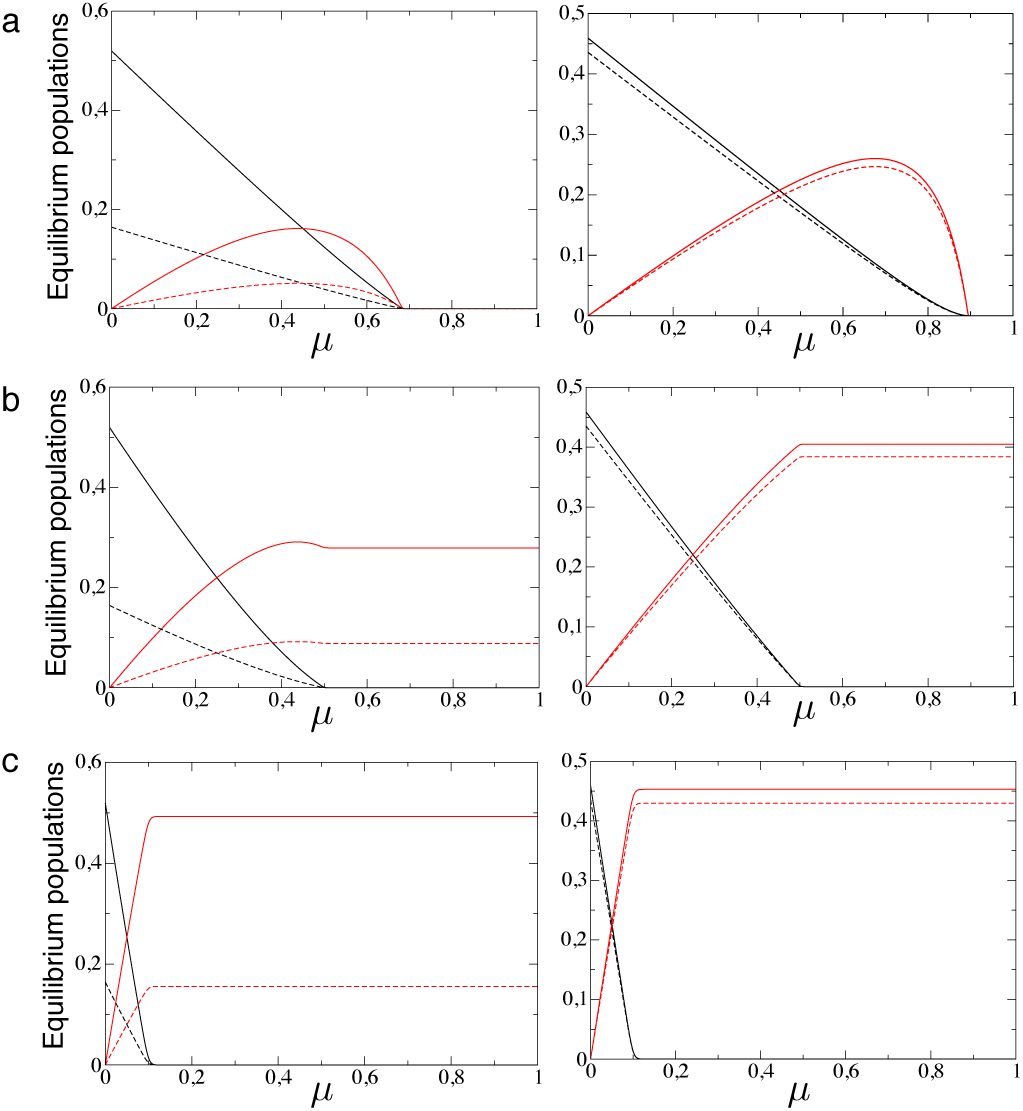
Equilibrium populations at increasing mutation rate *µ*, with *α* = 0.1 (first column) and *α* = 0.9 (second column). We analyze three different cases with: *k*_1_ = 0.1 (a); *k*_1_ = 0.5 (b); and *k*_1_ = 0.9 (c). In all of the panels we have set *ε* = 0.1 and the initial condition (*p*_0_(0),*n*_0_(0),*p*_1_(0),*n*_1_(0)) = (0.1, 0, 0, 0). Here, as in Fig. 4: (+) sense master (solid black line); (+) sense mutant (solid red line); (−) sense master (dashed black line); and (−) sense mutant (dashed red line).

In the following sections we generalize the results displayed in Figs. 4–6 by means of a deep analysis of the stability and the bifurcations of Eqs. (1)-(4).

**FIG. 6:**
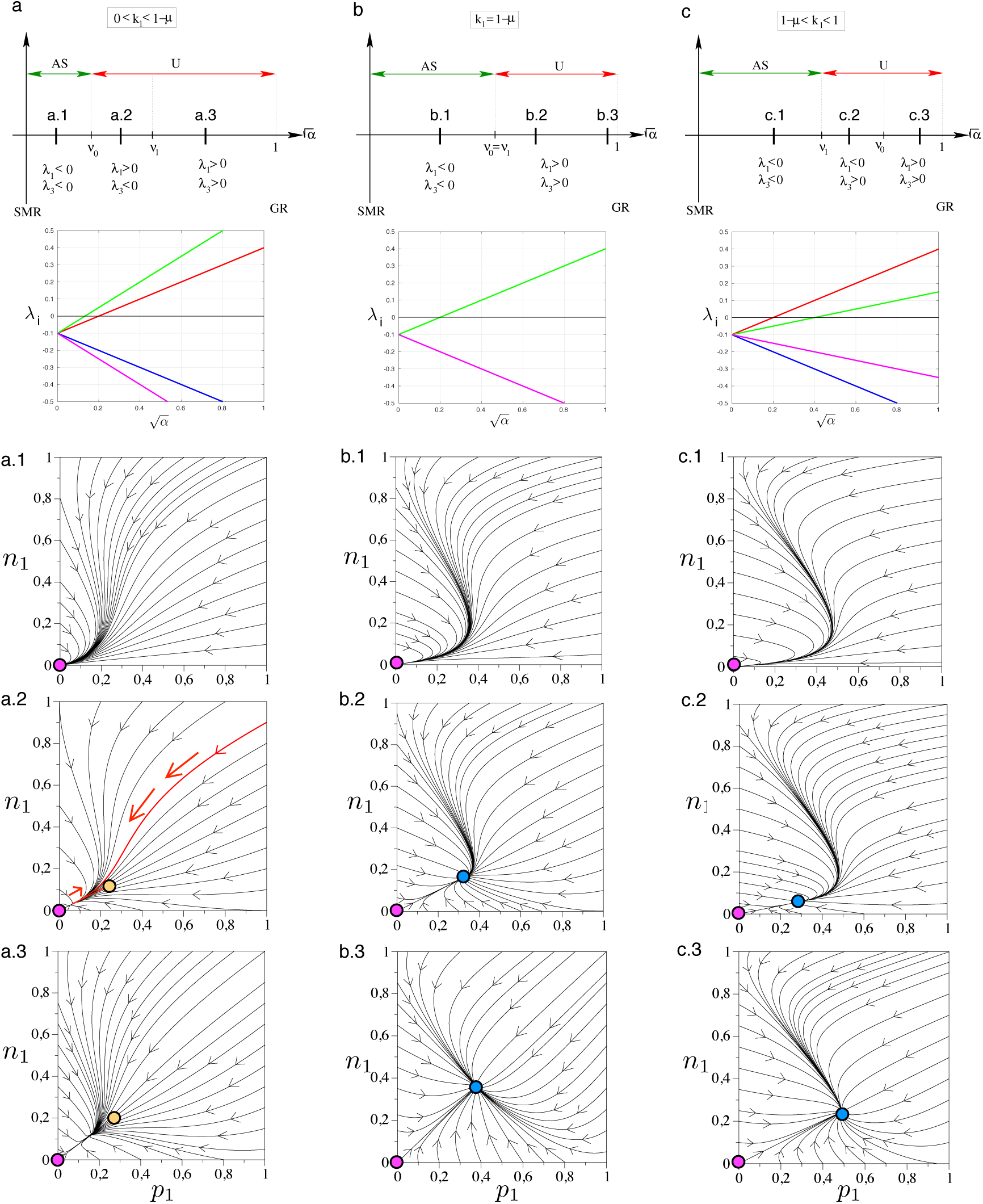
Local stability of the origin 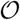 in three different scenarios: (a) 0 < *k*_1_ < 1 − *µ*; (b) *k*_1_ = 1 − *µ*; (c) *k*_1_ ≥ 1 − *µ* (AS means “asymptotically stable”; U denotes “unstable” and in all these cases means saddle type). Below each case we plot the eigenvalues of 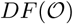 increasing 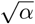 with *µ* = 0.5, *ϵ* = 0.1, and: *k*_1_ = 0.25 (a); *k*_1_ = 0.5 (b); and *k*_1_ = 0.75 (c). Here *λ*_1_ (red), *λ*_2_ (blue), *λ*_3_ (green), and *λ*_4_ (magenta). Phase portraits projected in the subspace (*p*_1_*,n*_1_) of the phase space Π are displayed setting *µ* = 0.6, *ϵ* = 0.1, and *k*_1_ = 0.15 (a), *k*_1_ = 0.4 (b), and *k*_1_ = 0.75 (c). Each panel corresponds to a value of 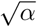: 0.15 (a.1); 0.25 (a.2); 0.75 (a.3); 0.15 (b.1); 0.5 (b.2); 0.95 (b.3); 0.09 (c.1); 0.2 (c.2); 0.5 (c.3). Fixed points: 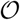 (magenta); 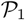 (blue); 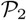 (orange). The red orbit in panel a.2 shows a trajectory that approaches the origin 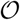 but then returns to 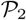.

## IV. LOCAL STABILITY OF THE EQUILIBRIA

This section is devoted to the study of the linear (and also in the majority of cases of the nonlinear) stability of the equilibria found in the previous section. We will consider separately the three equilibrium points 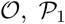 and 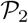. As it is standard, it will performed by considering the linearized system around them. Particular attention will be given to the change of stability of the equilibrium points that can indicate the presence of bifurcations, which are investigated in Section V. From now on we denote by *F* the vector field related to our system given by Eqs. (1)-(4).

### A. Stability of the origin

#### Proposition 3

*Let us consider the constants ν*_0_*,ν*_1_,*c_α_ defined in* (5)*. Then, the jacobian matrix at the origin DF* 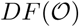 *has the following eigenvalues:*

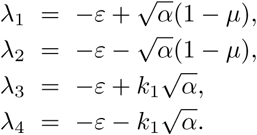

*Observe that all of them are real and that λ*_2_*,λ*_4_ *are always negative since* 0 < *µ* < 1 *and k*_1_ ≥ 0*. This means that the linear (and local nonlinear) stability of the origin will be determined by the signs of λ*_1_ *and λ*_3_*. Let us consider the following two cases:*

1. *Deleterious and neutral case* (0 < *k*_1_ ≤ 1)*: the three following scenarios hold:*

1. *If k*_1_ < 1 − *µ or, equivalently, ν*_0_ < *ν*_1_*: The origin* 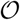 *is asymptotically stable (a sink) for* 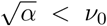 *and unstable for* 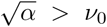. *For* 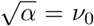 *we have the* birth *of* 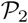*. More precisely, if* 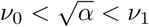 *then* dim 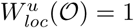 *and if* 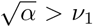 *then* dim 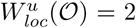, *where* 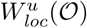 *denotes the local unstable invariant manifold of the equilibrium point* 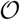.
2. *If k*_1_ = 1 − *µ or, equivalently, ν*_0_ = *ν*_1_ = *ν: In this situation*, 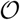 *is asymptotically stable (a sink) for* 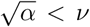 *and unstable for* 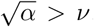*. This change in its stability coincides with the birth of* 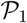*. Recall that if ν*_0_ = *ν*_1_ *the point* 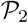 *does not exist. Moreover, when crossing the value* 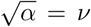 *one has that* dim 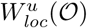 *passes from* 0 *to* 2.
3. *If k*_1_ > 1 − *µ or, equivalently, ν*_1_ < *ν*_0_*: Again, the origin is asymptotically stable (a sink) for* 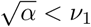 *and unstable for* 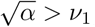, *coinciding with the birth of the equilibrium point* 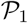*. As in the precedent case, no point* 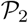 *exists. As above, if* 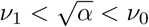 *then* dim 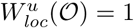 *and if* 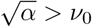 *then* 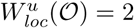,
2. *Lethal case* (*k*_1_ = 0)*: Taking into account again Proposition 1, the origin* 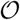 *changes its stability from asymptotically stable (a sink) to unstable (a saddle) when* 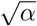 *crosses ν*_0_*. As above, this coincides with the birth of* 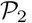.

Cases (i), (ii), and (iii) are displayed in Fig. 6a, b, and c, respectively. Specifically, the local stability of the origin for each case is shown as a function of 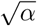: the upper panels in Fig. 6 display how the origin becomes unstable as the replication model changes from SMR to mixed modes. This means that under SMR the sequences are more prone to extinction, as suggested in [16]. These stability diagrams are also represented by means of the eigenvalues *λ*_1_…_4_. The phase portraits display the orbits in the subspace (*p*_1_*,n*_1_). Note that the label of each phase portrait corresponds to the letters in the upper panels. Panels a.1, b.1, and c.1 show results when the origin is a global attractor. Panels a.2 and a.3 display the orbits when the origin is unstable and the stable fixed point is 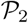, where the four genomes coexist. Finally, panels b.2, c.2, b.3, and c.3 display examples of a full dominance of the mutant genomes. For these latter examples, the increase of 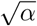 involves the change from the full extinction towards the survival of the mutant sequences.

### B. Stability of the point 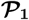

#### Proposition 4

*Let us assume* 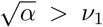, *in order the equilibrium points* 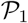 *to exist. Then, the eigenvalues of the jacobian matrix* 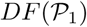 *are all real and they are given by*

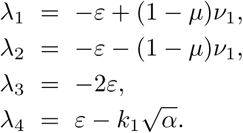

*The eigenvalues λ*_2_ *and λ*_3_ *are always negative. λ*_4_ *<* 0 *since* 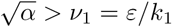*. Having in mind that ν*_0_ = *ε*/(1 − *µ*), *it is easy to check that:*

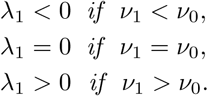

*Therefore, in the deleterious-neutral case we have the following subcases:*

1. *If k*_1_ < 1 − *µ or, equivalently, ν*_0_ < *ν*_1_: 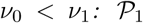 *is unstable (saddle). Indeed*, 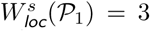 *and* 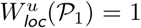, *where* 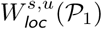 *denote the stable and unstable local invariant manifolds of* 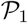.
2. *If k*_1_ = 1 − *µ or, equivalently, ν*_0_ = *ν*_1_ = *ν:* 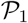 *has a* 1*-dimensional neutral direction (tangent to the eigenvector associated to the eigenvalue λ*_1_ = 0*) and a* 3*-dimensional local stable manifold*.
3. *If k*_1_ > 1 − *µ or, equivalently, ν*_1_ *<ν*_0_*: In this case* 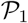 *is a sink so, therefore, a local attractor*.

*Regarding the lethal case* (*k*_1_ = 0), *the eigenvalue λ*_4_ = *ε is always positive and so* 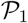 *is unstable (saddle)*.

The proof follows from straightforward computations.

Figure 7–8 provides the computation of the eigenvalues *λ*_1_…_4_ for the fixed point 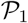.

**FIG. 7:**
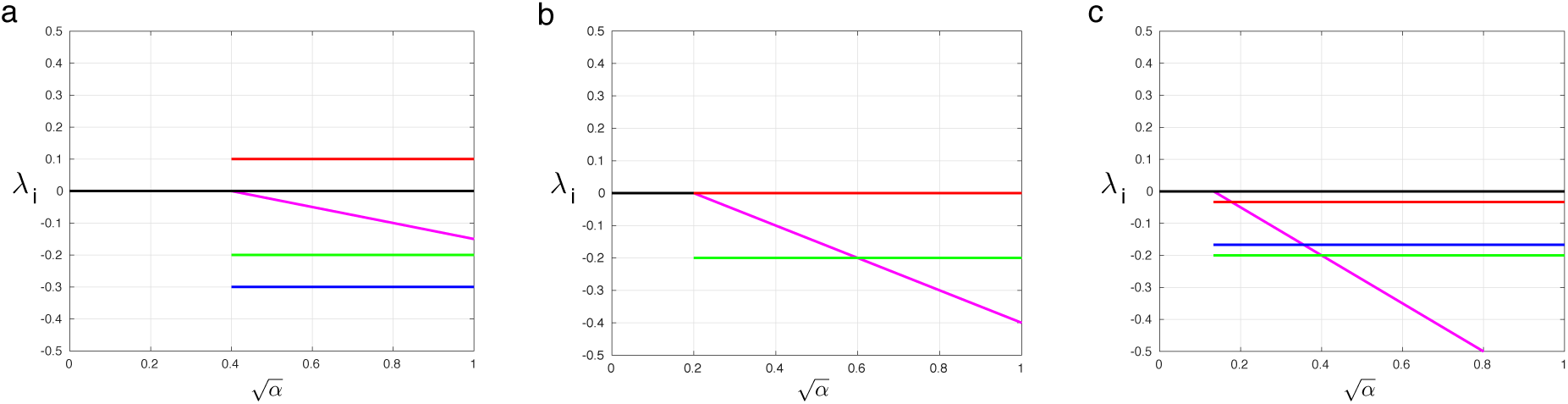
Eigenvalues of 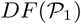 in the 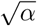 axis for the three different neutral-deleterious cases (i), (ii), and (iii) presented in Section IV.B. given by: *k*_1_ < 1 − *µ* (a); *k*_1_ = 1 − *µ* (b), and *k*_1_ > 1 − *µ* (c), respectively, with *µ* = 0.5 and *k*_1_ = 0.25, 0.5, and 0.75, respectively.: *λ*_1_ (red), *λ*_2_ (blue), *λ*_3_ (green) and *λ*_4_ (magenta). From left to right: The value of the parameter *ϵ* as been set to 0.1.

**FIG. 8:**
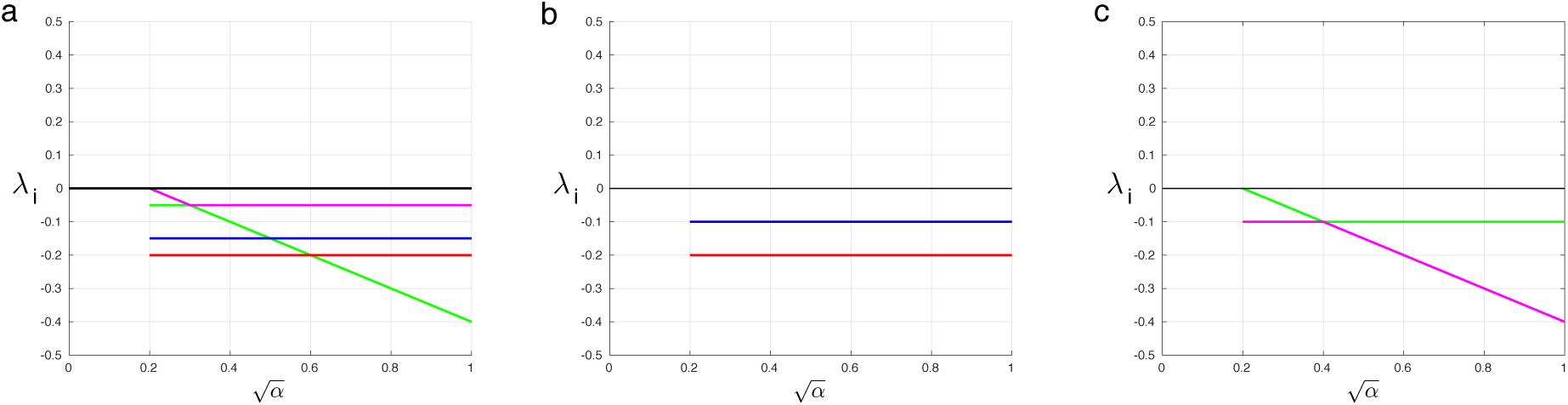
(a) Eigenvalues of 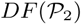 (deleterious case) in the 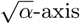 and 0 < *k*_1_ < 1 − *µ*: *λ*_1_ (red), *λ*_2_ (blue), *λ*_+_ (green) and *λ*_−_ (magenta). The parameters have been set to *µ* = 0.5, *k*_1_ = 0.25, and *ϵ* = 0.1. Remind that *ν*_0_ = *ε*/(1 − *µ*). (b-c) Eigenvalues of 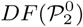 (lethal case) in the 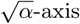: *λ*_1_ (red), *λ*_2_ (blue), *λ*_3_ (green) and *λ*_4_ (magenta). From left to right: (i) *λ*_1_,*λ*_2_; (ii) *λ*_+_,*λ*_−_ and (iii) all four eigenvalues with *µ* = 0.5 and *k*_1_ = 0.25. The value of the parameter *ϵ* as been set to 0.1.

### C. Stability of the points 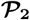 and 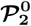

From Section III we know that the equilibrium point 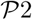 exists if 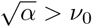 and in the following two cases:

1. In the deleterious case (0 < *k*_1_ < 1) provided that 0 < *k*_1_ < 1 − *µ* (or, equivalently, *ν*_0_ *<ν*_1_).
2. In the lethal case (*k*_1_ = 0).

Next proposition determines the local stability of 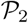 in these two situations.

#### Proposition 5

*Let us assume that* 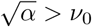 *in order* 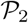 *and* 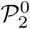 *to exist. Then, the eigenvalues of the differential* 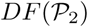 *and* 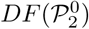 *are, respectively:*

1. *In the deleterious case (0 < k*_1_ < 1*) provided that 0* < *k*_1_ < 1 − *μ (or, equivalently, ν*_0_ < *ν*_1_):

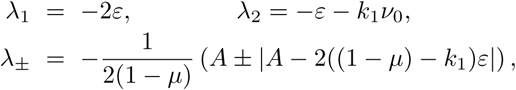
*where* 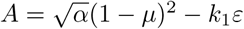*. Notice that assumptions* 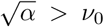 *and 0* < *k*_1_ < 1 − *μ imply that A* > 0.
2. *In the lethal case (k*_1_ = 0):

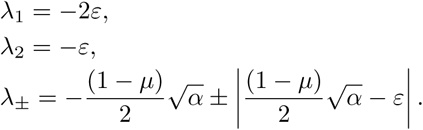 *Then, in both cases all four eigenvalues are real and negative, and so the equilibrium points* 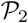 *and* 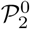 *are sinks for any* 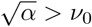.

Figure 8 displays the eigenvalues *λ*_1_…_4_ as a function of 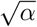 for the fixed point 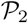 (Fig. 8a), and for the point 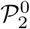 (Fig. 8b,c).

## V. BIFURCATIONS

Essentially the system suffers *transcritical* bifurcations. These bifurcations coincide with the appearance of a new equilibrium point, 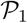, 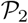 or 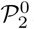. It is remarkable that the latter equilibria does not suffer, in principle, any bifurcation. Let us detail them in all our cases. Namely,

i. Deleteterious-neutral case (0 < *k*_1_ ≤ 1):

1. Case 0 < *k*_1_ < 1 − *µ* (that is, *ν*_0_ < *ν*_1_): the origin 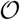 isa sink upto 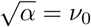. At that point, the equilibrium point 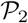 appears. Then, 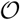 changes its stability by means of a transcritical bifurcation, becomes a saddle point (unstable), with 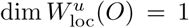. The coexistence equilibrium point 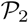 is a sink (i.e., an attractor) for 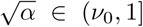. At 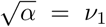, the equilibrium point 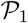 appears. It will be a saddle point (with 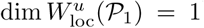) for 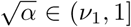. At this point, 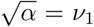, the dimension of 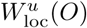 increases to 2, remaining like this up to 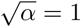.
2. Case *k*_1_ = 1 − *µ* (that is, *ν*_0_ = *ν*_1_): in this situation there are only two equilibrium points, 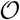 and 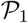, the latter appearing at 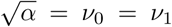. As above, the origin 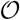 is a sink up to 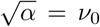. With the appearing of 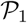 it undergoes a transcritical bifurcation leading it to unstable, precisely, a saddle point with 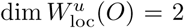. Concerning the point 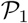, linearisation criterium does not decide its nonlinear local stability since it has (linear) centre and stable local invariant manifolds of dimension 1 and 3, respectively. No others bifurcations show up.
3. Case *k*_1_ > 1 − *µ* (that is, *ν*_1_ *<ν*_0_): similarly to the precedent cases, the origin is a sink (an attractor) until the appearance of the equilibrium 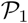 at 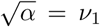. At this point, 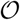 becomes unstable, a saddle with 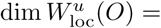. Later on, at 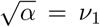, the dimension of 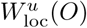 increases to 2, keeping this dimension until 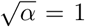. No bifurcations undergone by the point 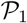, which is a sink for 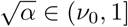.
ii. Lethal case (*k*_1_ = 0): there are only two equilibria: the origin 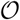 and the coexistence point 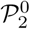, this latter appearing at 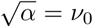. The origin is a sink for 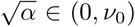, undergoes a transcritical bifurcation at 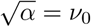, becoming unstable (saddle point) with 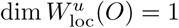. The point 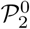 is always a sink.

Figure 9 summarizes the bifurcations found in Eqs. (1)-(4) obtained by choosing different values of *k*_1_ and tuning *α* from the SMR to the GR model. Here, for completeness, we overlap the information on stability for the origin, 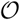, displayed in Fig. 6. Several phase portraits are displayed for each case. The panel in FIg. 9a shows the orbits for 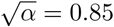 in the subspace (*p*_0_*,n*_0_), close to the GR mode. Here the attractor is 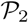, which is asymptotically globally stable and involves the coexistence between master and mutant genomes. For the case *k*_1_ = 1− *µ* and for 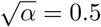 the attractor achieved is 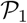, indicating that the population is dominated by the pool of mutants at equilibrium (Fig. 9b). The same asymptotic dynamics is found in the phase portrait of Fig. 9c. Finally, for *k*_1_ = 0 we plot a case for which 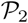 is also globally asymptotically stabe, while 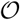 is unstable (Fig. 9d).

**FIG. 9:**
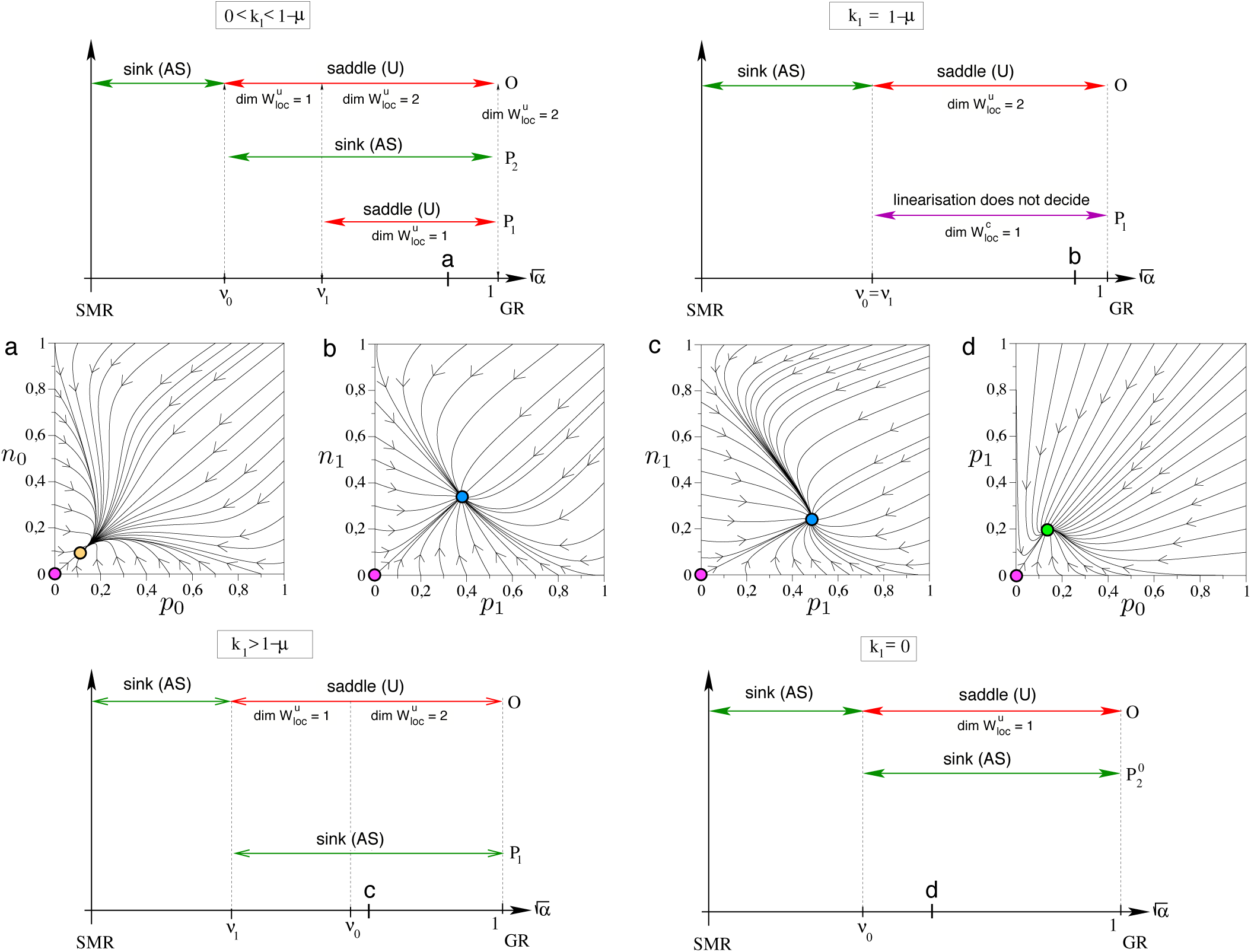
Bifurcations of the equilibrium points 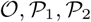 (deleterious-neutral cases) and 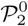 (lethal case). From top to bottom and left to right: deleterious-neutral case, (i) 0 < *k*_1_ < 1 − *µ*, (ii) *k*_1_ = 1 − *µ*, (iii) *k*_1_ > 1 − *µ*; and (iv) lethal case. The phase portraits correspond to the parameter values indicated with the letters in the bifurcation diagrams with: *k*_1_ and 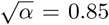 (a); *k*_1_ = 0.4 and 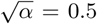 (b); *k*_1_ = 0.75 and 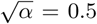; and *k*_1_ = 0, 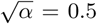 (b). Initial conditions: *p*_1_(0) = *n*_1_(0) = 0 (a); *p*_0_(0) = *n*_0_(0) = 0.1 (b); and *p*_0_(0) = *n*_0_(0) = 0 (c-d). In all of the panels we use *µ* = 0.6 and *ε* = 0.1. Fixed points: 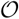 (magenta); 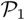 (blue); 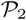 (orange); 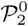 (green).

Let us now focus our attention on the bifurcation diagram for the deleterious-neutral case. In this context, for a given value 0 < *µ* < 1 we consider a plane in the parameters 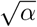 and *k*_1_. By hypothesis (H2), the diagram is restricted to the rectangle 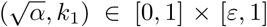. The bifurcation curves 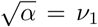 and 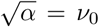 are, respectively, the hyperbola 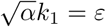 and the vertical line 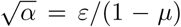. The three colored areas in Fig. 10a correspond to the *ω*-limit of the solution starting with initial conditions *p*_0_(0) = 1, *n*_0_(0) = *p*_1_(0) = *n*_1_(0) = 0 (the same result hold with *p*_0_(0) = 0.1, *n*_0_(0) = *p*_1_(0) = *n*_1_(0) = 0). Namely, convergenc to the origin 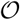 (red area); convergence to the equilibrium point 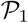 (light green area); attraction by the equilibrium point *P*_2_ (blue area). Observe that, when crossing these two bifurcation curves the equilibrium points change stability - by means of a transcritical bifurcation - or change the dimension of its associated local unstable invariant manifold (when they are saddles).

**FIG. 10:**
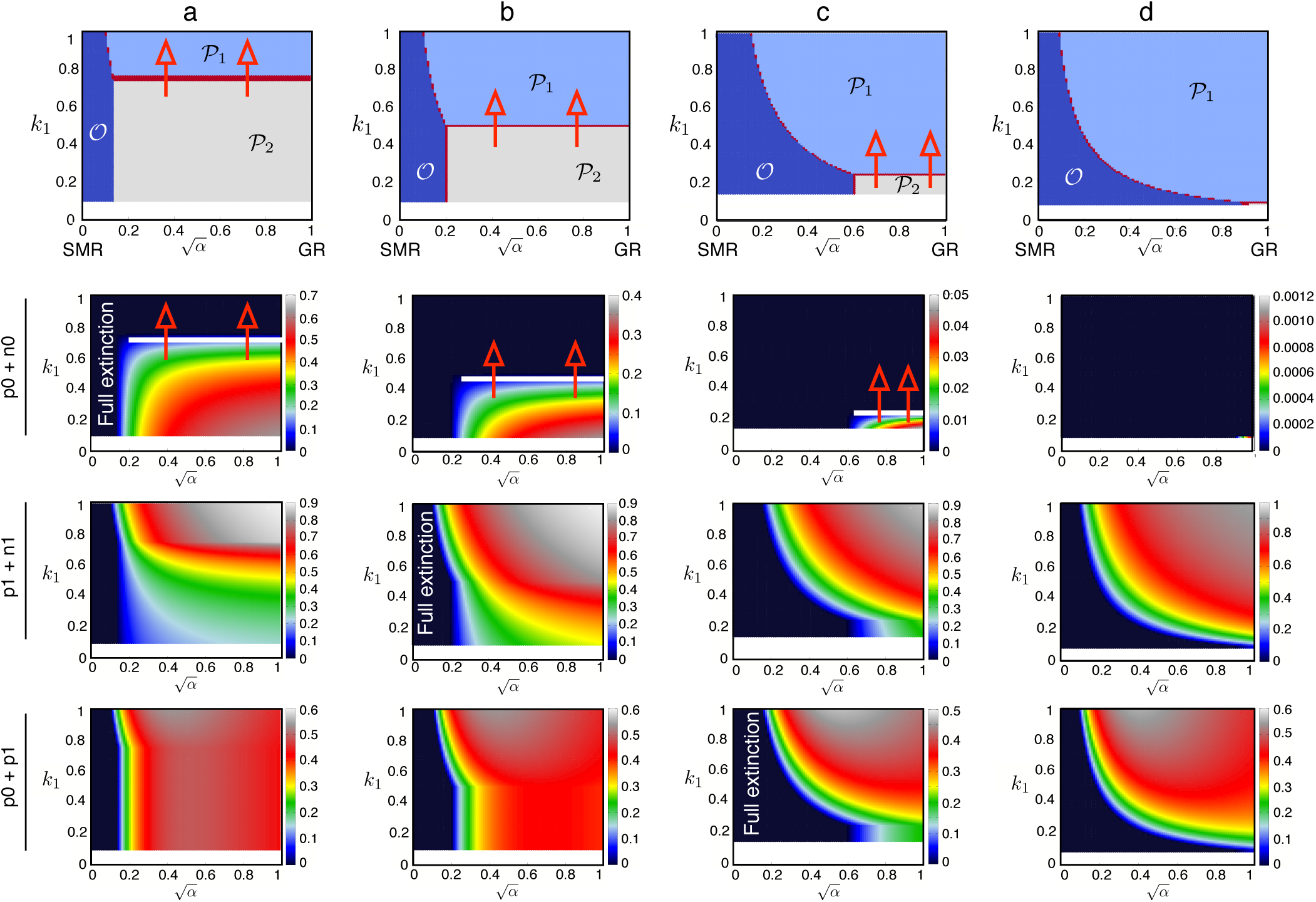
Two-dimensional parameter spaces displaying the stability of the fixed points. (a) 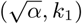-plane bifurcation diagram for the deleterious-neutral cases. The thick red line indicates the boundary for the full dominance of the mutant sequences as a function of *k*_1_. Crossing this boundary (vertical red arrows) causes the extinction of the master sequences *p*_0_*,n*_0_ and the dominance of the pool of mutants (green surface). Below this line all genomes coexist (blue area). (b) 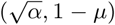-plane bifurcation diagram indicating the stability of the fixed points for the lethal case. The vertical black lines indicate the entry into lethal mutagenesis, where full extinctions occur (light blue). The regions with survival of all sequences is colored in orange.

Similarly, we can plot a bifurcation diagram in the lethal case (*k*_1_ = 0, Fig. 10b), now depending on the parameters 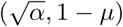. Again, hypothesis (H2) implies that it takes places in the rectangle [0, 1] × [*ε*, 1]. The bifurcation curve 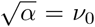 becomes a branch of the hyperbola 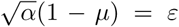. This curve also divides the domain in two coloured areas: a blue one, at the left-hand side of the hyperbola, characterized by the fact that the equilibrium point 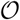, the origin, is the *ω*-limit of the solution starting at the initial conditions *p*_0_(0) = 1, *n*_0_(0) = *p*_1_(0) = *n*_1_(0) = 0; an orange one, located on the right-hand side of the hyperbola, where the equilibrium point 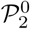 is this *ω*-limit. Figure 11 displays the regions in the parameter space 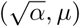 where the different asymptotic states (obtained numerically) can be found for the detelerious-neutral cases: sequences extinction (red); dominance of mutant sequences (green); and coexistence of sequences (blue). Notice that these regions obtained numerically perfectly match with the analytical results derived in the article. In this plot we can identify the critical mutation values causing lethal mutagenesis (yellow arrows in Fig. 11), which occurs for 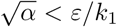. Above this threshold, lethal mutagenesis is replaced by the error catastrophe (red line in Fig. 11), with a critical mutation rate not depending on *α*. Notice that when the replication mode is close to the SMR lethal mutagenesis is achieved for lower mutation rates. This means that replication modes departing from the SMR provide the sequences with more resistance to lethal mutagenesis.

**FIG. 11:**
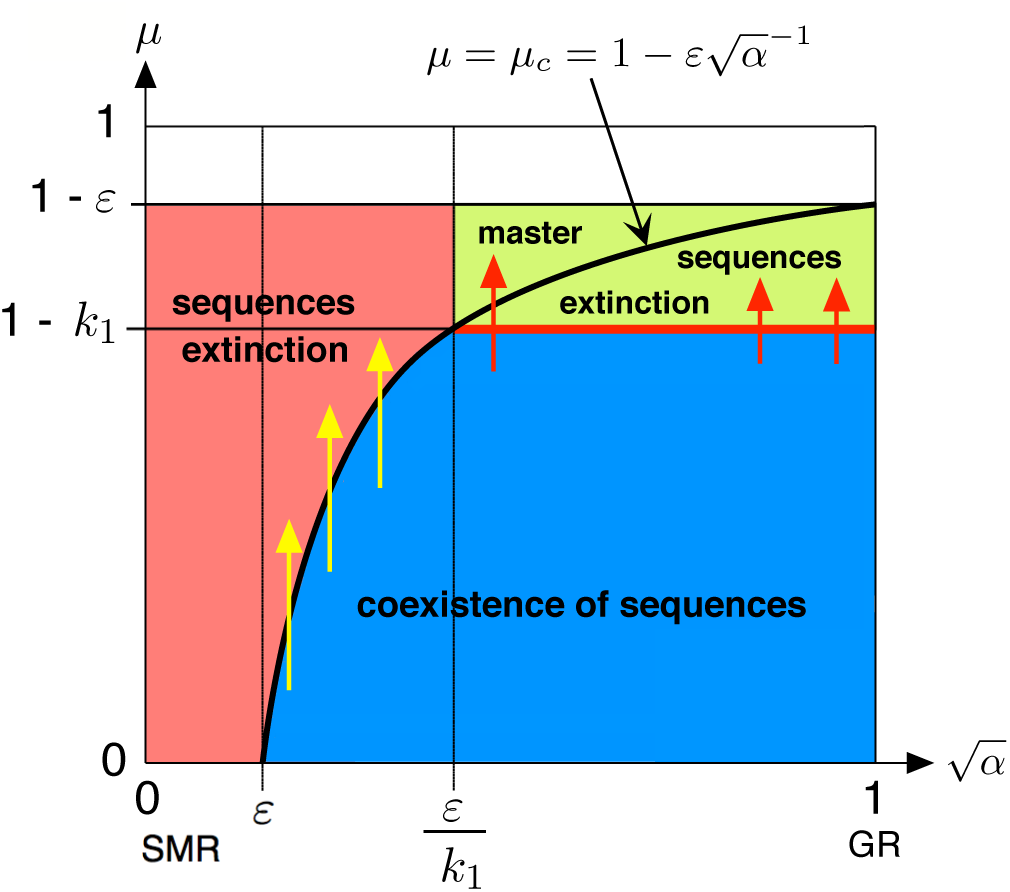
Phase diagrams for the deleterious-neutral case computed numerically in the parameter space 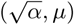. The equilibrium state is represented using the same colors than in Fig. 10a. The critical mutation rates involving the entrance into error threshold is displayed in red. The yellow arrows indicate the entrance into lethal mutagenesis. This plot has been built using (*p*_0_(0) = 0.1,*n*_0_(0) = 0*,p*_1_(0) = 0*,n*_1_(0) = 0) as initial conditions. The same results are obtained with initial conditions (1, 0, 0, 0). Notice that lethal mutagenesis is replaced by the error catastrophe as *α* increases.

Finally, in Fig. 12 we display the basins of attraction of the fixed points for the neutral and deleterious mutants displayed in Fig. 10a. The red arrows indicate those values of *k*_1_ responsible for the dominance of the mutant sequences (first and second rows in Fig. 12). Also, we computed numerically the relative populations for the master genomes (second row in Fig. 12), as well as of the mutants (third row) and the master and mutant (+) sense sequences.

**FIG. 12:**
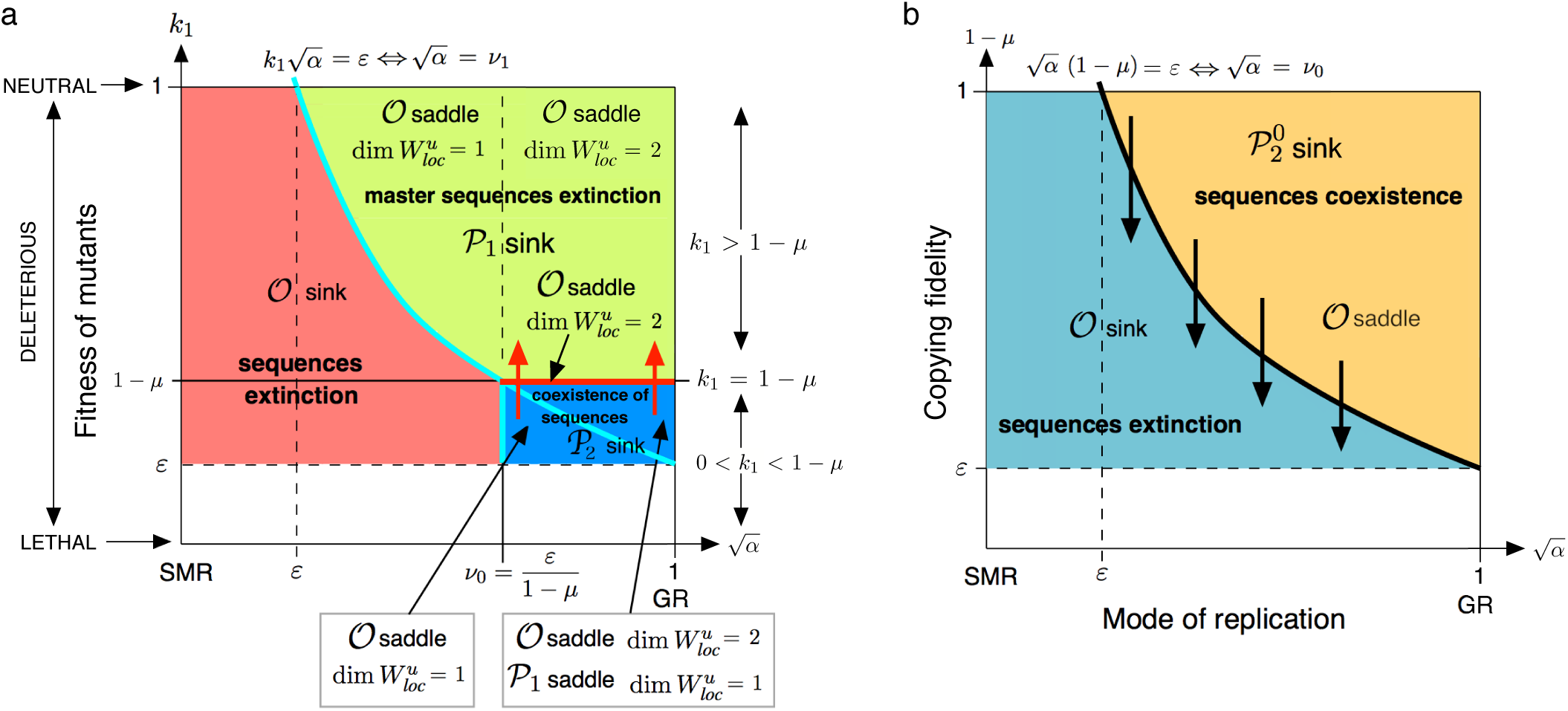
Phase diagrams for the deleterious-neutral case displayed in Fig. 10a. We display the asymptotic dynamics in the parameter space 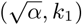, with (a) *µ* = 0.25 and *ε* = 0.1; (b) *µ* = 0.5 and *ε* = 0.1; (c) *µ* = 0.75 and *ε* = 0.15; (d) *µ* = 0.9 and *ε* = 0.09. Legend: origin 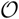 (dark blue); 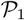 (light-blue); 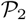 (light-grey); 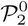 (light-red) and “no convergence” (dark red). Below the phase diagrams we display the equilibrium populations obtained numerically for variables: *p*_0_ + *n*_0_ (upper row); *p*_1_ + *n*_1_ (mid row); and *p*_0_ + *p*_1_ (lower row) 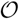. The horizontal white lines in the upper row display those critical values *k*_1_ involving the dominance of the mutant sequences.

## VI. CONCLUSIONS

The evolutionary dynamics of RNA viruses has been largely investigated seeking for critical thresholds involving error catastrophes and lethal mutagenesis [17, 18, 22]. Previous research on viral RNA replication modes has focused on theoretical and computational studies aiming at describing the evolutionary outcome of RNA sequences under the Stamping Machine Replication (SMR) and Geometric Replication (GR) modes. Smooth transitions have been identified [13, 19]. For instance, a simple model considering (+) and (−) sense genomes under differential replication modes identified a transcritical bifurcation [16]. This model, however, did not consider evolution. In this article we have studied a simple model considering both (+) and (−) sense sequences with differential replication modes and evolving on a single-peak fitness landscape. Despite the simplicity of this landscape, being highly unrealistic, it has been used in multiple models as a simple approach to the dynamics of RNA viruses [17, 18]. The model studied here has allowed us to derive the critical mutation values involving error thresholds and lethal mutagenesis considering three different types of mutants spectra, given by neutral, deleterious, and lethal mutants.

In the deleterious case, there are three possible scenarios when increasing the value of *µ* (we omit the trivial total extinction solution which is always assumed as a possible equilibrium): if 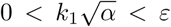, that is, close to the SMR mode, there is no nontrivial equilibrium solution. This happens for any *µ* > 0. In the region of parameters 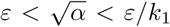, between the SMR and GR modes (depending on the particular values of *ε* and *k*_1_), the bifurcation undergone by the equilibria is quite steep. It passes from a situation with coexistence equilibrium to total extinction equilibrium when crossing the curve 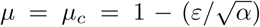. For 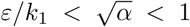, which includes always the GR case. When increasing *µ*, the systems evolves from coexistence to master sequences’ extinction when crossing the critical value *µ* = 1 − *k*_1_.

Summarizing, the error threshold is achieved when the mutation rate is above the critical value *µ_c_*, in the deleterious case is given by 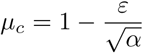 if 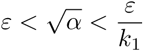; and *µ_c_* = 1 − *k*_1_ if 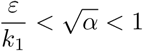. In the lethal case, there are only two scenarios: for 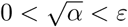 (that is, almost pure SMR-mode), there are no nontrivial equilibria. For the rest of cases, that is, 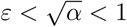 the possible equilibrium solution goes from coexistence to total extinction. Our results have allowed us to relate the processes of lethal mutagenesis and error catastrophe, establishing the parametric conditions making theoretical viral quasispecies to shift from one phenomenon to the other.

## Acknowledgements

The research leading to these results has received funding from “la Caixa” Foundation and from a MINECO grant awarded to the Barcelona Graduate School of Mathematics (BGSMath) under the ”María de Maeztu” Program (grant MDM-2014-0445). JS and TA has been also partially funded by the CERCA Programe of the Generalitat de Catalunya. JTL has been partially supported by the MINECO/FEDER grant MTM2015-65715-P, by the Catalan grant 2014SGR-504 and by the Russian Scientific Foundation grants 14-41-00044 and 14-12-00811. TA is also supported by the AGAUR (grant 2014SGR-1307), the MINECO (grant MTM2015-71509-C2-1-R). SFE has been supported by MINECO-FEDER grant BFU2015-65037-P and by Generalitat Valenciana grant PROMETEOII/2014/021.

## VII. APPENDIX

### A. Proof of Proposition 1

Let us deal, first, with the deleterious case (0 < *k*_1_ < 1). In this framework, equilibrium states will come from the solutions of the following system of non-linear equations:

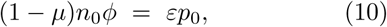

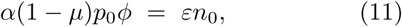

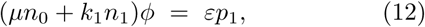

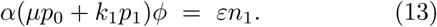

It is clear that the origin 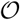 is a fixed point of our system in all the cases. To find nontrivial solutions we distinguish three different scenarios for these equilibria: (i) master sequences extinction; (ii) mutant sequences extinction and (iii) coexistence among all sequences.

(i) Case *p*_0_ = *n*_0_ = 0 (master sequences extinction): If we assume *p*_1_ = 0, substituting in equation (13) and using that *ε* ≠ 0, we get *n*_1_ = 0 and there fore, the equilibrium is 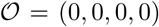, the trivial solution. A symmetric situation undergoes when we start taking *n*_1_ = 0. Thus, let us assume that *p*_1_*n*_1_ ≠ 0. Replacing *p*_0_ = *n*_0_ = 0 in (12)–(13) and dividing such equations we get *p*_1_/*n*_1_ = *n*_1_/(*αp*_1_) and so 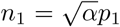. This division is well-defined since *p*_1_ > 0, *k*_1_ > 0 and *ϕ* ≠ 0 (if *ϕ* = 0 it is straightforward to check that it leads to the origin 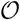 as fixed point). From equation (13) we obtain 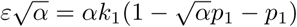 and thus

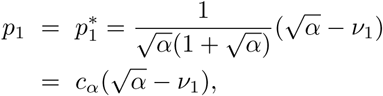
where *ν*_1_ and *c_α_* have been defined in (5). Therefore, since 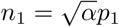 we get the equilibrium point 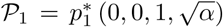 provided 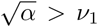 (since we are interested in nontrivial equilibrium points with biological meaning).
(ii) Case *p*_1_ = *n*_1_ = 0 (mutant sequence extinction): in this scenario one has to solve

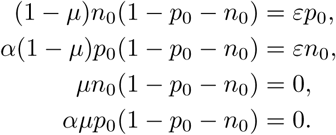 As before, both cases *p*_0_ = 0 and *n*_0_ = 0 lead to the equilibrium point 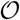. So let us consider the case of *p*_0_*n*_0_ ≠ 0. From the last two equations it follows that *p*_0_ + *n*_0_ = 1 and substituting in the two ones we get *p*_0_ = *n*_0_ = 0, which is a contradiction. So there is no nontrivial equilibrium points with *p*_1_ = *n*_1_ = 0.
(iii) Coexistence sequences equilibria: multiplying equation (11) by *p*_0_ and subtracting equation (10) multiplied by *n*_0_ it turns out that 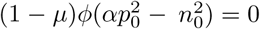. Since 0 < *µ* < 1, this leads to three possibilities, namely, (a) *ϕ* = 0 (that is *p*_0_+*n*_0_+*p*_1_+*n*_1_ = 1) or (b) 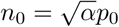 with *ϕ* ≠ 0 and (c) *ϕ* = 0 and 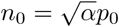. Case (c) does not apply. Indeed, substituting *ϕ* = 0 and 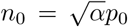 into equation (10) one gets that *p*_0_ = 0 and so *n*_0_ = 0, which is not possible. A similar argument shows that case (a) does not happen. In fact, taking *ϕ* = 0 in equations (10)–(13) leads to *p*_0_ = *n*_0_ = *p*_1_ = *n*_1_ = 0 which contradicts *ϕ* = 0 ⇔ *p*_0_ + *n*_0_ + *p*_1_ + *n*_1_ = 1. Thus, let us deal with case (b). Substituting 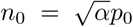 in (10) and using that *p*_0_ ≠ 0 (if *p*_0_ = 0 ⇒ *n*_0_ = 0, which corresponds to the master sequences extinction case) it turns out that

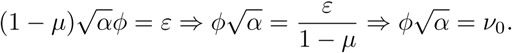

It is straightforward to check that equation (11) leads to the same condition. Performing again the change 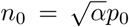 onto equations (12) and (13) one gets

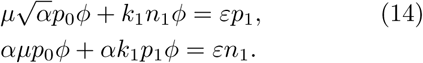

Computing the division between equation (12) and (13), namely,

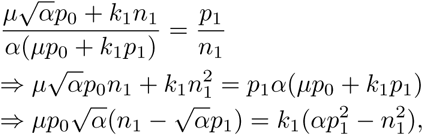
one gets

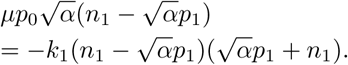

So now we have two possibilities: 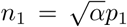 or 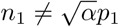. Observe that the latter cannot be since in that case we would have that 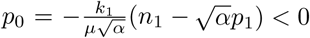, which is not possible because *p*_0_ is positive. Therefore, it must be 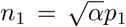. Substituting it into (14) we have *µp*_0_*ν*_0_ + *k*_1_*p*_1_*ν*_0_ = *εp*_1_, which implies

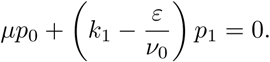

Notice that *k*_1_ − (*ε*/*ν*_0_) = 0 ⇔ *ν*_0_ = *ν*_1_. In fact, we have that *ν*_0_ ≠ *ν*_1_. Indeed, if this term vanished we would have *p*_0_ = 0 and thus *n*_0_ = 0, which gives rise to point 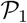.

Hence, if *k*_1_ − (*ε*/*ν*_0_) ≠ 0, it follows that

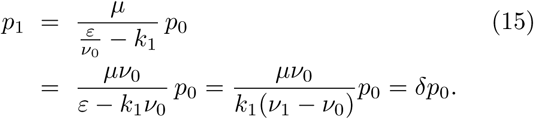

Thus, 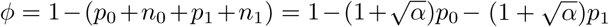 and so

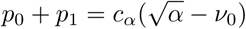

Combining the previous relation with (15) the following solution is obtained

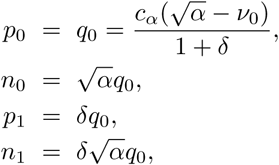
with *ν*_0_, *c_α_*, *δ* defined in (5)–(6), which leads to the coexistence equilibrium state

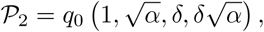
for 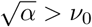 and *ν*_0_ < *ν*_1_.

Concerning the neutral case (*k*_1_ = 1), it is easy to check that all the computations carried out for the deleterious context are also valid for this case.

And the last, but not least, case corresponds to the lethal framework (*k*_1_ = 0). Equilibrium states must be solution of the system

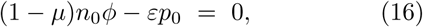

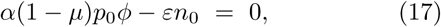

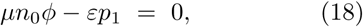

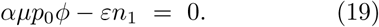

Again, the origin 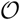 is a trivial fixed point. To seek for nontrivial equilibria we take into account two scenarios: (a) *p*_0_ = 0; (b) *p*_0_ ≠ 0.

1. Case *p*_0_ = 0: From the equation (16) we get (1 − *µ*)*n*_0_*ϕ* = 0. Since 0 < *µ* < 1 we have three possibilities: *n*_0_ = 0, *ϕ* = 0 or both. It is obvious that first and third cases lead to the origin 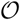. Regarding to the case with *ϕ* = 0, it follows that *n*_0_ + *n*_1_ + *p*_1_ = 1. Substituting it into equations (17)–(19) we get *n*_0_ = *p*_1_ = *n*_1_ = 0, which contradicts the previous equality.
2. Case *p*_0_ ≠ 0: From (16) we have that neither *n*_0_ nor *ϕ* vanish. Performing *n*_0_ × (16) minus *p*_0_ × (17) one gets that 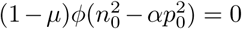 and so 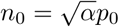 since 0 < *µ* < 1 and *ϕ* ≠ 0. Substituting the latter equality into (16) it follows that 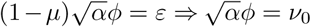. Subtracting *n*_0_ × (19) from *αp*_0_ × (18) one has 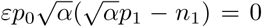, so then 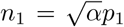. On the other hand, 

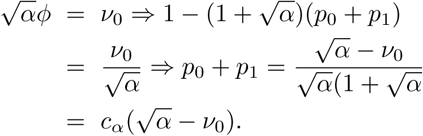 And last, from (19) and using that 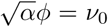 and 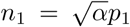 we get 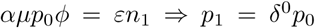. Therefore the equilibrium point is given by

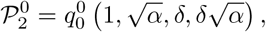
where 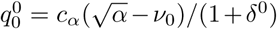 and provided that 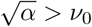 (to have biological meaning).

#### B. Proof of Proposition 2

As mentioned before, the case *µ* = 1 corresponds to the situation when there is no autocatalysis in the master sequence and so it mutates with probability 1. Thus, concerning their equilibrium points we have:

- In the deleterious and neutral cases, substituting *µ* = 1 into equations (10)–(13), one gets the equations

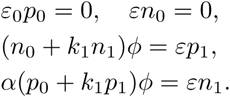 From the two first equations it follows that *p*_0_ = *n*_0_ = 0 and, consequently

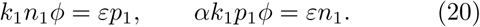 Again, we distinguish several possibilities:

- If *n*_1_ = 0 then *p*_1_ = 0 and so we obtain the origin.
- If *p*_1_ = 0 then *n*_1_ = 0 and therefore the equilibrium point is again the origin.
- In case that *n*_1_ + *p*_1_ = 1, *n*_1_ ≠ 0, *p*_1_ ≠ 0 it follows that *ϕ* = 0 and so *p*_1_ = *n*_1_ = 0 which is a contradiction with the fact that *n*_1_ + *p*_1_ = 1.
- Finally, if *n*_1_ ≠ 0, *p*_1_ ≠ 0, *ϕ* ≠ 0, we can divide them and get *αp*_1_/*n*_1_ = *n*_1_/*p*_1_. Consequently, 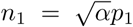. This gives rise to an equilibrium of the form 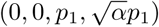. Substituting this form into the first equation of (20), one obtains 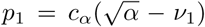, defined provided 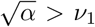, which corresponds to the point 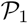 in Proposition 1.
- In the lethal case, equilibria system (16)-(19) reduces to *εp*_0_ = 0, *εn*_0_ = 0, *n*_0_*ϕ* = *εp*_1_, *αp*_0_*ϕ* = *εn*_1_. From the first two equations we have *p*_0_ = *n*_0_ = 0 and substituting in the second ones, it turns out *p*_1_ = *n*_1_ = 0, that is, the origin.

#### C. Proof of Proposition 3

As usual, we use stability analysis of the linearised system around the equilibrium to determine, when possible, the local nonlinear stability of the point for the complete system.

1. Deleterious and neutral case (0 < *k*_1_ ≤ 1): the eigenvalues of the differential matrix

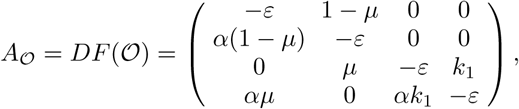
are 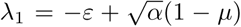, 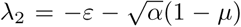, 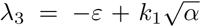, and 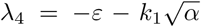. It is easy to verify that 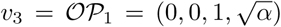 and 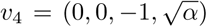 are eigenvectors of *λ*_3_ and *λ*_4_, respectively. It is also straightforward to check that

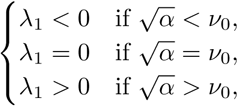
and

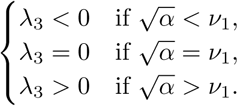

Thus, we have the following three cases: can check that

- Case 0 < *k*_1_ < 1 − *µ* or, equivalently, *ν*_0_ < *ν*_1_: the origin is a sink (an attractor) for *α* ∈ (0*,ν*_0_) and unstable (saddle) for 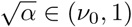. For *α* ∈ (*ν*_0_*,ν*_1_) one has 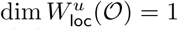 and if 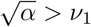 then 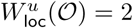.
- Case *k*_1_ = 1 − *µ* or, equivalently, *ν*_0_ = *ν*_1_: the origin is a sink for 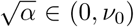 and unstable (saddle) for 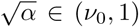. The dimension of 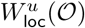 is 2 in this interval.
- Case 1 − *µ* < *k*_1_ < 1 or, equivalently, *ν*_1_ > *ν*_0_: the origin is a sink if 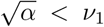 and unstable (a saddle) for 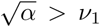. The dimension 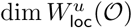 goes from 1 to 2 when 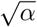 crosses *ν*_0_.

2. Lethal case (k1 = 0): The eigenvalues of

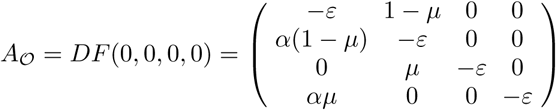
are in this case

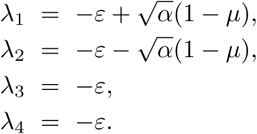

Observe that *λ*_2_ < 0, *λ*_3_ < 0 and *λ*_4_ < 0 so the stability of 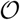 depends only on *λ*_1_. Indeed:

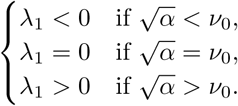

Therefore, the origin is asymptotically stable for 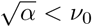 and becomes unstable for 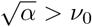. This situation is represented in Fig. 6.

#### D. Proof of Proposition 5

Remind that 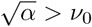 since 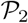 exists. We distinguish two cases:

1. Case 1: deleterious mutants (0 < *k*_1_ < 1) with 0 < *k*_1_ < 1 − *µ* (that is, equivalently, *ν*_0_ < *ν*_1_). The expression of the eigenvalues can directly from algebraic computations. They are all real. Observe that *λ*_1_, *λ*_2_ and *λ*_+_ are negative. Concerning *λ*_−_, notice that

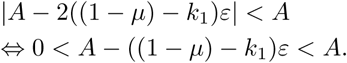

The second inequality is trivially satisfied since (1− *µ*) − *k*_1_ > 0 and *ε* > 0. Regarding the first one, one can check that

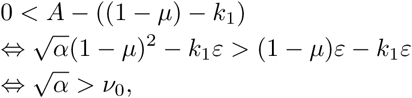
which is satisfied by hypothesis. Therefore, *A*−|*A*−2((1 − *μ*) − *k*_1_)*ε*| > 0 and, consequently, λ_−_ < 0. This implies that the point 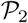 is a sink for any 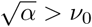.

2. Case 2: lethal mutants (*k*_1_ = 0). As above, the expression for the eigenvalues follows from linear algebra and straightforward computations. Again, *λ*_1_*,λ*_2_, and *λ*_−_ are all three real and negatives. Concerning *λ*_+_ (real), we define 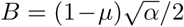. This implies that *λ*_+_ = −*B* + |*B* − *ε*|. Observe that |*B* − *ε*| < *B* ⇔ 0 < 2*B* − *ε*. Right-hand inequality is trivial since *ε* > 0. Left-hand is also satisfied since it is equivalent to 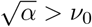. So, all four eigenvalues are real and negative which means that the point 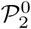 is a sink for any 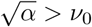.

## References

[1] R. Sanjuán, M.R. Nebot, N. Chirico, L.M. Mansky, and R. Belshaw. Viral mutation rates. J. Virol., 84:9733–9748, 2010.

[2] R. Sanjuán and P. Domingo-Calap. Mechanisms of viral mutation. Cell Mol. Life Sci., 73:4433–3338, 2016.

[3] M. Eigen. Self organization of matter and the evolution of biological macromolecules. Naturwissenschaften, 58(10):465–523, 1971.

[4] G. Stent. Molecular Biology of Bacterial Viruses. 1963.

[5] S. Luria. The frequency distribution of spontaneous bacteriophage mutants as evidence for the exponential rate of phage reproduction. Cold Spring Harbor Symp. Quant. Biol., 16:463–470, 1951.

[6] L. Chao, C.U. Rang, and L.E. Wong. Distribution of spontaneous mutants and inferences about the replication mode of the rna bacteriophage *ϕ*6. J. Virol., 76:3276–3281, 2002.

[7] L. Garcia-Villada and J.W. Drake. The three faces of riboviral spontaneous mutation: spectrum, mode of genome replication, and mutation rate. PLoS Genet., 8:e1002832, 2012.

[8] F. Martínez, J. Sardanyès, S.F. Elena, and J.A. Daròs. Dynamics of a plant rna virus intracellular accumulation: stamping machine vs. geometric replication. Genetics, 188:637–646, 2011.

[9] M.B. Schulte, J.A. Draghi, J.B. Plotkin, and Andino R. Experimentally guided models reveal replication principles that shape the mutation distribution of rna viruses. eLife, 4:e03753, 2015.

[10] M. Combe, R. Garijo, R. Geller, J.M. Cuevas, and R. Sanjuán. Single-cell analysis of rna virus infection identifies multiple genetically diverse viral genomes within single infectious units. Cell Host Microbe, 18.

[11] C.A. III Hutchison and R.L. Sinsheimer. The process of infection with bacteriophage *ϕx*174. x. mutations in a *ϕx* lysis gene. J. Mol. Biol., 18:429–447, 1966.

[12] S.F. Elena, P. Carrasco, J.A. Daròs, and R. Sanjuán. Mechanisms of genetic robustness in rna viruses. EMBO Rep., 7:168–173, 2006.

[13] J. Sardanyés, R.V. Solé, and Elena S.F. Replication mode and landscape topology differentially affect rna virus mutational load and robustness. J. Virol., 83:12579–12589, 2010.

[14] J. Sardanyés and S.F. Elena. Quasispecies spatial models for RNA viruses with different replication modes and infection strategies. PloS one, 6(9):e24884, 2011.

[15] J. Sardanyés, R.V. Solé, and S.F. Elena. Replication mode and landscape topology differentially affect RNA virus mutational load and robustness. J. Virol., 83(23):12579–89, 2009.

[16] F. Sardanyès, J. Martínez, J.A. Daròs, and S.F. Elena. Dynamics of alternative modes of rna replication for positive-sense rna viruses. J. Roy. Soc. Interface, 11:768–776, 2012.

[17] R. Pastor-Satorras and R.V. Solé. Field theory for a reaction-diffusion model of quasispecies dynamics. Physical Review E, 64(5):051909, 2001.

[18] R.V. Solé, J. Sardanyés, J. Díez, and A. Mas. Information catastrophe in RNA viruses through replication thresholds. Journal of Theoretical Biology, 240(3):353–359, 2006.

[19] J.J. Bull, L.A. Meyers, and M. Lachmann. Quasispecies made simple. PLoS computational biology, 1(6):e61, 2005.

[20] R. Sanjuán, A. Moya, and S.F. Elena. The distribution of fitness effects caused by single-nucleotide substitutions in an rna virus. Proc. Natl. Acad. Sci. U.S.A., 101:8396–8401, 2004.

[21] P. Carrasco, F. de la Iglesia, and S.F. Elena. Distribution of fitness and virulence effects caused by single-nucleotide substitutions in tobacco etch virus. J. Virol., 81:12979–12984, 2007.

[22] S.C. Manrubia, E. Domingo, and E. Lázaro. Pathways to extinction: beyond the error threshold. Philos. Trans. R. Soc. Lond. B, 365:1943–1952, 2010.

